# Peritoneal macrophage phenotype correlates with pain scores in women with suspected endometriosis

**DOI:** 10.1101/2020.07.31.209106

**Authors:** Douglas A Gibson, Frances Collins, Bianca De Leo, Andrew W Horne, Philippa TK Saunders

## Abstract

**Objective:** To characterise peritoneal macrophage populations in women with suspected endometriosis and assess if they are correlated with severity of pelvic pain symptoms.

**Design:** Flow cytometry analysis of peritoneal fluid samples and clinical data.

**Setting:** University Research Institute.

**Patients:** Clinical questionnaires, surgical data and peritoneal fluid were collected with informed consent from women undergoing diagnostic laparoscopy for suspected endometriosis (n=54).

**Intervention(s):** None

**Main Outcome Measure(s):** Severity of pelvic pain symptoms was assessed by the EHP-30 questionnaire. Immune cells recovered from peritoneal fluid were analysed by flow cytometry.

**Results:** Pain scores (pain domain of EHP30) did not differ according to endometriosis diagnosis, stage of endometriosis or whether or not women were receiving hormone treatment. Analysis of immune cells in peritoneal fluid revealed two populations of peritoneal macrophages: CD14^high^ and CD14^low^ which were not altered by menstrual cycle phase or hormone treatment. CD14^high^ peritoneal macrophages were increased in women with endometriosis compared to those without but were not altered by coincident reproductive health issues such as infertility or heavy menstrual bleeding. Peritoneal macrophage phenotype correlated with pelvic pain symptoms in women with suspected endometriosis. Notably, CD14^high^ peritoneal macrophages negatively correlated with pain scores whereas CD14^low^ peritoneal macrophages were positively correlated. This association was independent of endometriosis diagnosis.

**Conclusion:** Peritoneal macrophage phenotypes correlate with pelvic pain symptoms in women with suspected endometriosis and are altered by presence of disease. These results provide new insight into the association between endometriosis pathophysiology and pelvic pain symptoms.

## Introduction

Endometriosis is a chronic neuroinflammatory pain condition affecting ∼180 million women worldwide and has a socioeconomic impact similar to diabetes (1). It is defined by the presence of endometrial-like tissue outside the uterus (‘lesions’), commonly within the pelvic cavity. The most common symptoms of endometriosis are chronic pelvic pain, dyspareunia, and dysmenorrhoea (1, 2). While the exact aetiology of endometriosis is unknown, changes in inflammatory processes are thought to contribute to the pathogenesis of the disease.

Peritoneal macrophages are the most abundant immune cell in the peritoneal cavity (3) with defined roles in clearance of apoptotic cells and immune surveillance. Studies in mouse models suggest that peritoneal macrophages are required for the establishment and persistence of endometriosis lesions. For example, the number of peritoneal macrophages is increased in mice with experimentally-induced endometriosis compared to controls (4) and macrophages expressing the angiopoietin receptor Tie-2 are reported to accumulate in lesions and promote lesion vascularisation and growth (5). Depletion of peritoneal macrophages prevents lesion formation and depleting peritoneal macrophages in mice with established lesions reduces lesion vascularisation and prevents further growth (5).

These mechanistic studies in mice have been complemented by studies comparing peritoneal fluid from women with and without endometriosis. Notably, increased concentrations of macrophage growth factors and chemokines such as colony stimulating factor 1 (CSF1) and monocyte chemoattractant protein 1 (CCL2) are detected in peritoneal fluid from women with endometriosis (6-8) and this is associated with increased numbers of peritoneal macrophages compared to women without disease (8-10). Furthermore, peritoneal macrophage function appears to be dysregulated in endometriosis patients and this is characterised by increased secretion of pro-inflammatory cytokines and reduced phagocytic capacity (8, 11-13). Thus, peritoneal macrophage dysfunction may contribute to persistence of endometrial tissue and establishment of lesions in the cavity.

Macrophage dysfunction may also affect pain symptoms via secretion of neurotrophic factors that promote nerve growth (14). Notably, CD68-positive macrophages co-localise with nerve fibres within peritoneal endometriotic lesions and it has been speculated that they can promote endometriosis pain symptoms in women (15). It is not clear if peritoneal macrophages directly contribute to pain symptoms in women but in a mouse model of endometriosis peritoneal macrophages appear to promote endometriosis-associated hyperalgesia, putatively via secretion of macrophage-derived insulin-like growth factor 1 (16).

To date, peritoneal macrophages have largely been assessed as a single population in women with endometriosis and this has limited our understanding of their function. However, recent studies in other disorders have reported there is heterogeneity within human peritoneal macrophage populations based on expression of complement receptor of the immunoglobulin superfamily (CRIg) (17) or canonical markers, such as CD14, CD16 and HLA-DR (18, 19). Notably, peritoneal macrophage subsets appear to be altered in some diseases, such as liver cirrhosis (17). It is not known whether different subtypes of peritoneal macrophage are present in endometriosis, and if so, whether their phenotype is associated with disease severity or pain symptoms.

The objective of this study was to phenotype and quantify peritoneal macrophage subpopulations in the peritoneal fluid of women with suspected endometriosis and determine if specific populations are associated with endometriosis stage, symptoms, or hormone treatment.

## Material and methods

### Study approval

Written informed consent was obtained from all study participants prior to surgery; ethical approval was granted by the Lothian Research Ethics Committee (LREC 11/AL/0376). Methods were carried out in accordance with NHS Lothian Tissue Governance guidelines and EPHect guidelines (https://endometriosisfoundation.org/ephect/).

### Clinical samples and patient data

Peritoneal fluid was recovered from women undergoing diagnostic laparoscopy for suspected endometriosis in NHS Lothian (n=54). Pain scores were determined using the pain domain of the Endometriosis Health Profile Questionnaire (EHP-30) (a validated endometriosis-specific pain questionnaire (20)). Age, BMI, menstrual cycle stage and hormone status were obtained and recorded along with other key clinical data (Supplementary Table 1 and 2).

For women who were not on hormones and had regular menstrual cycles, the diagnosis of endometriosis was confirmed macroscopically at laparoscopy in nineteen women, whereas no evidence of endometriosis was found in ten women (‘no endo’). Women with endometriosis diagnosis were subsequently classified according to the revised American Fertility Society guidelines (rAFS) as stage I (n=10), II (n=4) and III/IV (n=5). For some analyses, women who were receiving hormone treatment were included (n=25). In women who met this criterion, diagnosis of endometriosis was confirmed in fourteen women, while eleven were classified as ‘no endo’ (no obvious pelvic pathology at laparoscopy).

### Flow Cytometry

Immune cells recovered from peritoneal fluid were analysed by flow cytometry using standard methods (21). Briefly, cells were washed and subjected to red cell lysis before being counted and resuspended in FACS buffer at a concentration of 0.5-1×10^6^ cells/100µL. Antibodies were used at manufacturer’s recommended concentration (Supplementary table 3). Cells were gated to select immune cells (live, CD45^+^) and further sub-gated to select myeloid cells (CD11b^+^) and exclude T, B and NK cells (CD3^-^, CD19^-^, CD56^-^). Gating of subpopulations was performed based on fluorescence minus one (FMO) and population distribution (CD14). This population gating was applied to all samples. Samples were assessed using a Becton Dickinson FACS Aria II. Data were analysed using FlowJo™ Software.

### Statistics

Statistical analysis was performed using Graphpad prism. T test was used to test the difference in the means of two groups. Two-way ANOVA was used to determine significance between treatments in grouped data. Non-parametric testing was utilised where sample sizes were insufficient to confirm normality of data distribution; Kruskal-Wallis test was used to assess differences between multiple groups or Mann Whitney test to assess variance between two groups. Pearson correlation was used to quantify the degree to which two variables (macrophages abundance and pain score) were related. Criterion for significance was p<0.05. All data are presented as mean ± SEM.

## Results

### Pelvic pain does not correlate with endometriosis diagnosis or AFS stage

Baseline characteristics of the patient cohort were assessed to identify potential factors which may influence pelvic pain in women with suspected endometriosis. Pain scores were determined using the pain domain of the EHP-30 questionnaire (20). For women who were not on hormones, Age, BMI, and pain scores did not differ between those women categorised as ‘no endo’ or those with surgically diagnosed endometriosis (Fig.1A and Supplementary table 1). There was no statistically significant difference in pain score between AFS stages I, II or III/IV, although pain scores were highly variable among some AFS stages (Fig. 1B). Oral contraceptives and treatments which suppress hormone production are reported to decrease endometriosis-associated pain in some women (22, 23). In this study, pain scores in women receiving hormonal treatment (H) were not different to those women who were not on hormones (NH) and this was similar in women with (Fig. 1C) and without (Fig. 1D) a confirmed surgical diagnosis of endometriosis.

**Figure 1.**
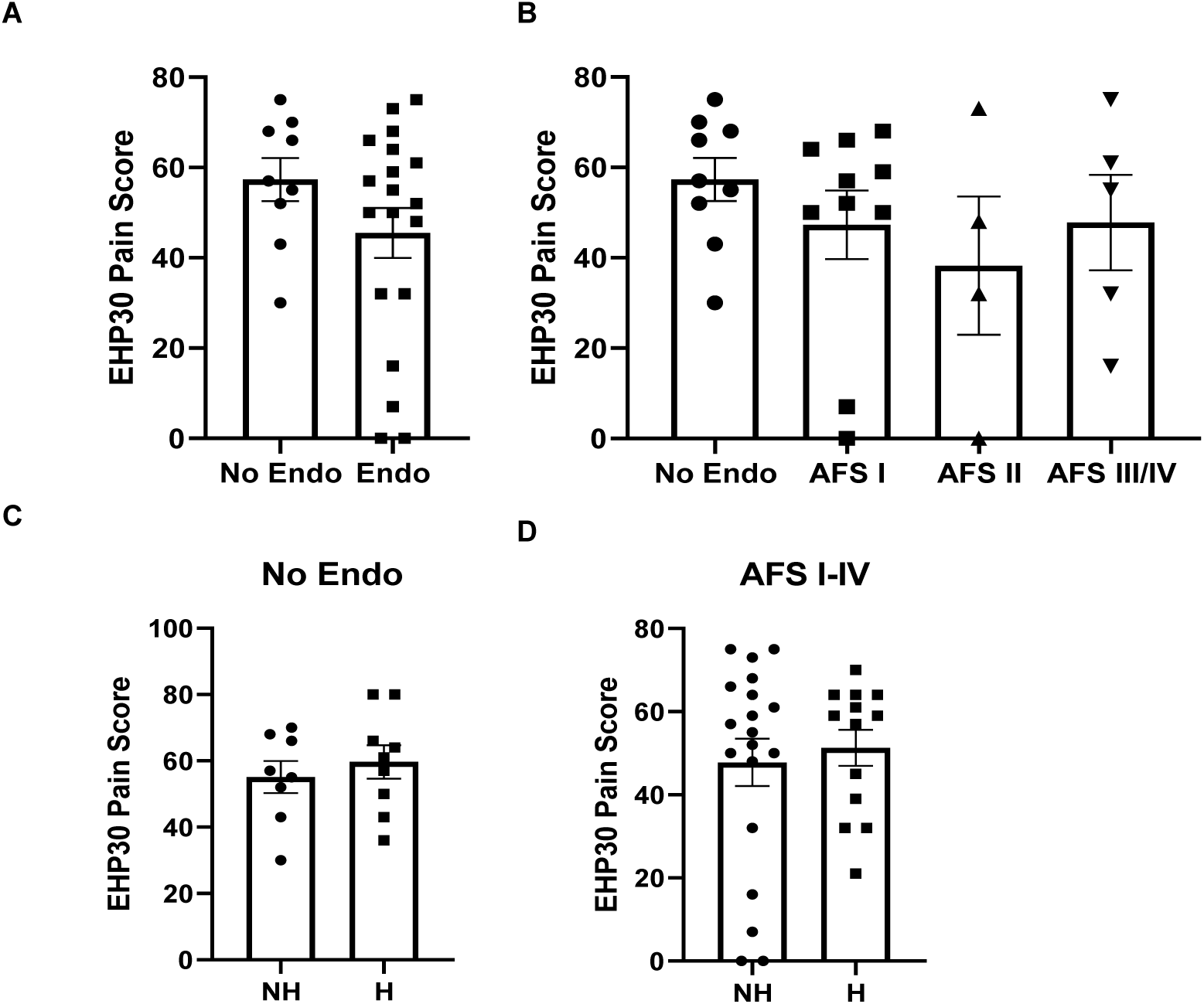
Pain score does not differ by endometriosis diagnosis, stage of disease or hormone treatment. **A** Presence/absence of endometriosis did not affect pain score (pain domain of EHP30) and **B** did not differ according to AFS stage. Hormone treatment did affect pain score in women without (**C)** or with endometriosis diagnosis **(D)**.

### Two populations of peritoneal macrophages are present in the peritoneal fluid of women with suspected endometriosis and they are not altered by hormone status

Cells were recovered from peritoneal fluid and analysed by flow cytometry. Cells were gated to select immune cells and further sub-gated to select myeloid cells and exclude T, B and NK cells (Fig. 2A &B). Two populations of peritoneal macrophages (PMφ) were identified which were HLA-DR-positive and had either high or low intensity expression of CD14 (Fig. 2C).

**Figure 2.**
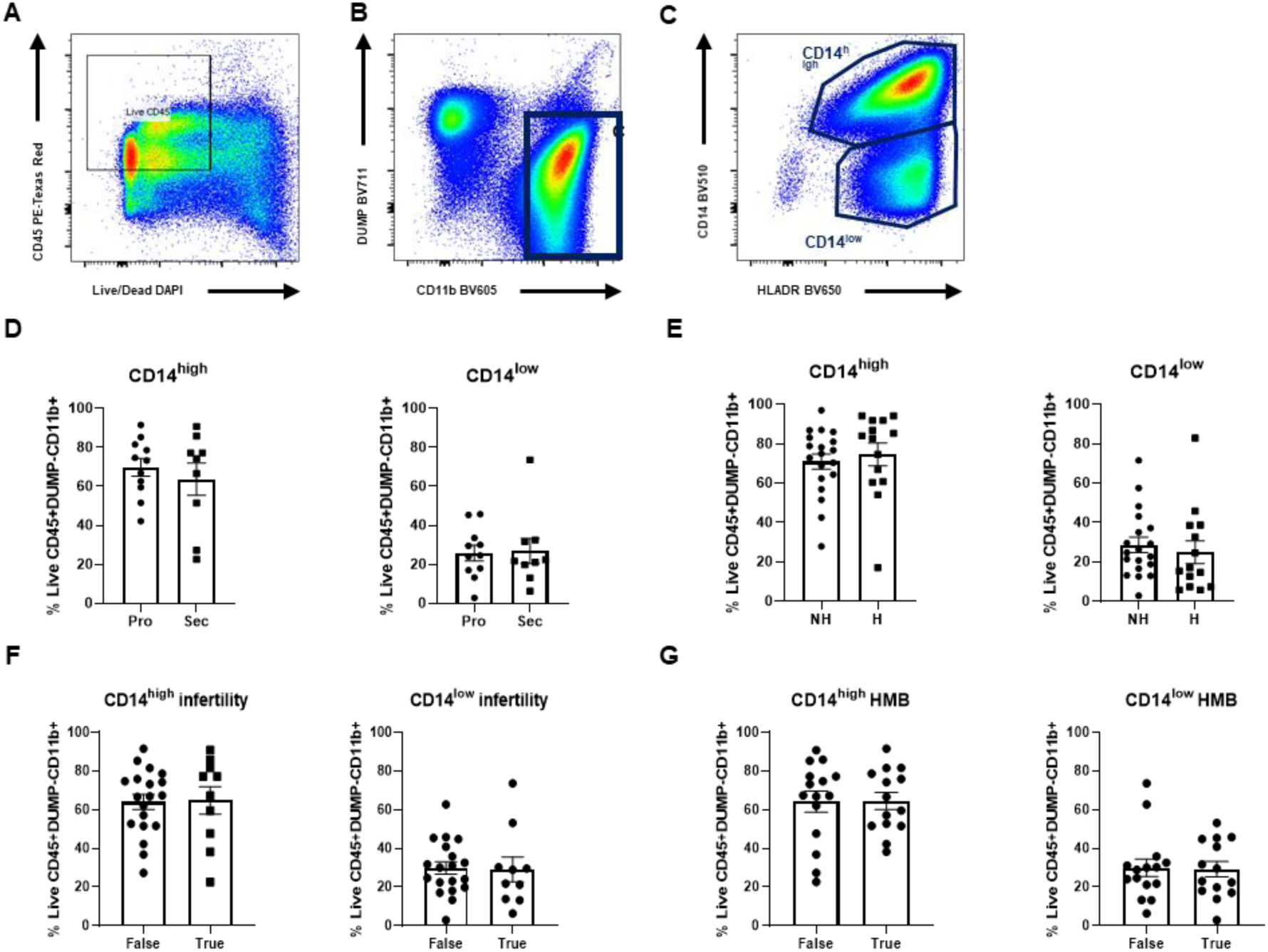
Analysis of peritoneal fluid immune cells from women with diagnosed endometriosis reveals two populations of peritoneal macrophages that are not altered by hormones or other reproductive health disorders. Representative flow cytometry plots from pelvic peritoneal fluid (PF) immune cells. Cells were gated to select; **A** Live, CD45^+^ (all immune cells) and **B** CD11b^+^ (myeloid cells), CD3^-^CD19^-^ CD56^-^ (‘DUMP’; T, B and NK cells). **C** Two peritoneal macrophage (PMφ) subpopulations were identified that were HLA-DR^+^ and CD14^+^ (High vs low). **D** Phase of menstrual cycle (Pro: proliferative; Sec: secretory) did not affect the abundance of CD14^high^ (*t* test, p=0.5035) or CD14^low^ cells (Mann Whitney test, p=0.7664) thus samples were thereafter not stratified on this basis. **E** The abundance of CD14^high^ (Mann Whitney test, p=0.3723) or CD14^low^ cells (Mann Whitney test, p=0.3345) did not differ if women were not on hormones (NH) or on a hormonal treatment (H). **F** The abundance of CD14^high^ or CD14^low^ peritoneal macrophages (PMφ) did not differ between women reporting infertility (True) and those who did not (False). **G** The abundance of CD14^high^ or CD14^low^ PMφ in women with endometriosis did not differ between women reporting HMB (True) and those who did not (False).

In women with regular menstrual cycles, the phase of the menstrual cycle (categorised as proliferative or secretory), had no significant impact on the abundance of CD14^high^ or CD14^low^ PMφ populations (Fig. 2D). Although hormone treatment did not alter abundance of any of these immune cell populations (Fig. 2E; supplementary figure 1), samples from patients on hormones were excluded from subsequent analysis to avoid any confounding effects of this variable on other disease parameters.

### Peritoneal macrophages are not altered by infertility or heavy menstrual bleeding

We next assessed whether the co-morbidities infertility or heavy menstrual bleeding (HMB), recorded as part of patient cohort data collection, could affect the abundance of PMφ subpopulations in women with endometriosis. Infertility was more common in women with endometriosis than women without endometriosis (42% vs 20%; supplementary table 1), however, infertility did not alter the abundance of CD14^high^ or CD14^low^ macrophages in women with endometriosis (Fig. 2F). HMB was less common in women with endometriosis than in women without endometriosis (42% vs 60%; supplementary table 1) but PMφ populations were not altered by presence of HMB (Fig. 2G).

### CD14^high^ peritoneal macrophages are increased in women with endometriosis

PMφ were assessed in women with and without endometriosis who had regular cycles and were not receiving hormone treatment (‘No endo’ vs ‘AFSI-IV’). CD14^high^ PMφ were significantly higher in the women with endometriosis (AFS I-IV) compared to women without disease (Fig. 3A, p<0.05). No difference was observed for CD14^low^ PMφ, although the average was lower for women with endometriosis (Fig. 3B). To assess the relationship between the two populations we next assessed the ratio of CD14^high^ to CD14^low^ macrophages. CD14^high^ PMφ were the predominant population in the peritoneal cavity and this difference was more pronounced in women with endometriosis (Fig. 3C) and with increasing AFS stage (Fig. 3D). Analysis according to AFS stage (Figure 3C) identified a significant difference between subpopulations of CD14^high^ and CD14^low^ PMφ in women classified as AFS stage I (p<0.01), II (p<0.0001) or III/IV (p<0.001). No significant difference was found between CD14^high^ and CD14^low^ PMφ in women without endometriosis.

**Figure 3.**
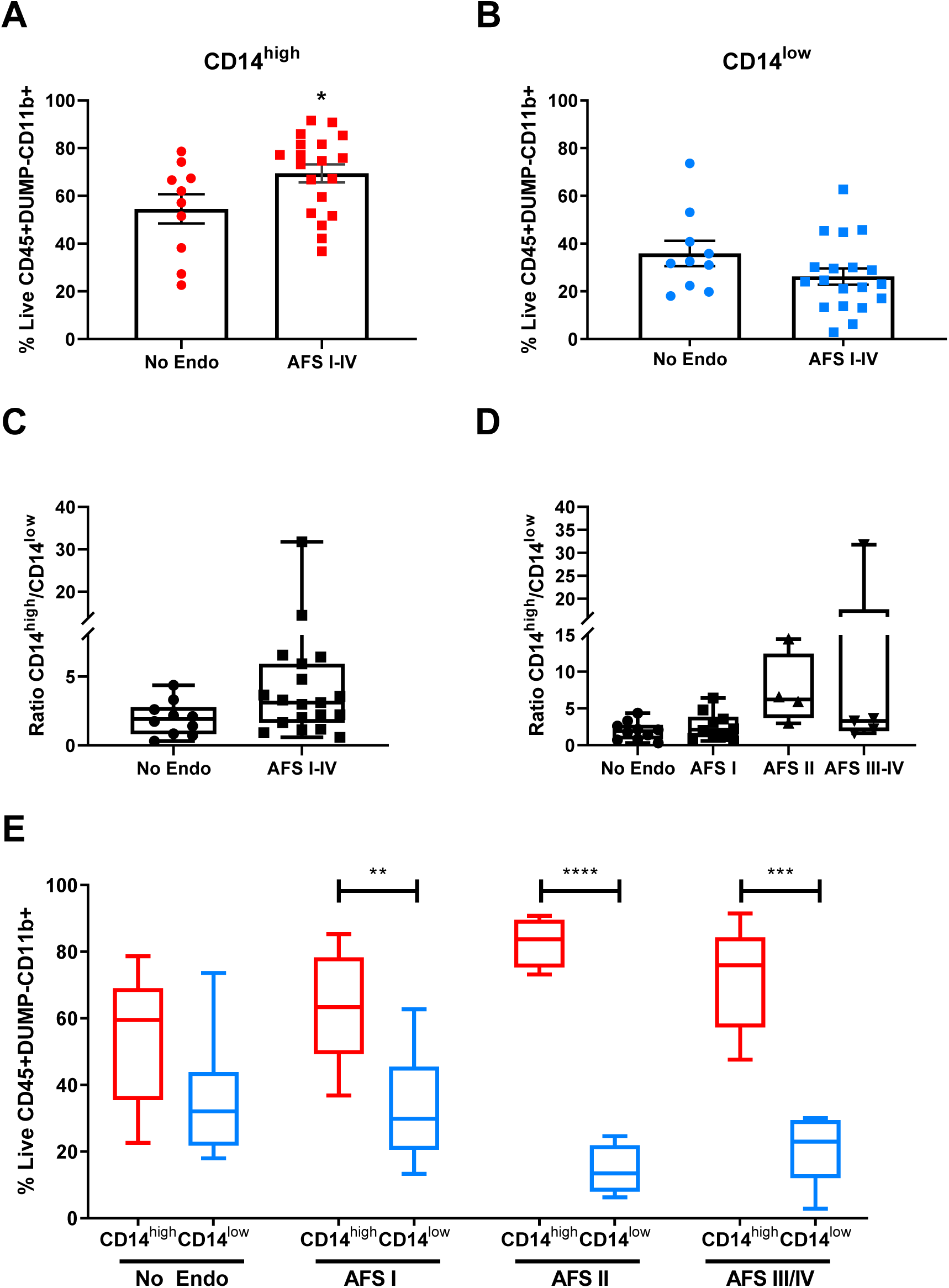
CD14^high^ peritoneal macrophages are increased in women with endometriosis but not altered by other. **A** Frequency of CD14^high^ peritoneal macrophages (PMφ) was greater in women with endometriosis (AFS stage I-IV, n=19, t test, p<0.05) than those without endometriosis (‘No endo’; n=10). **B** Frequency of CD14^low^ PMφ was not significantly different in women with endometriosis compared to those without disease **C** The ratio of CD14^high^ to CD14^low^ PMφ was greater in women with endometriosis (AFSI-IV) than those without endometriosis (no endo). **D** The ratio of CD14^high^ to CD14^low^ PMφ was greater in women from each AFS stage than those without endometriosis (no endo). **E** CD14^high^ PMφ were significantly higher than CD14^low^ PMφ in women with endometriosis stratified according to AFS I (n=10, P<0.01), AFS II (n=4, P>0.0001) and AFS III/IV (n=5 P>0.001) but not in women without endometriosis (n=9; Two-way ANOVA). *p<0.05, **p<0.01, ***p<0.001, ****p<0.0001.

### Peritoneal macrophages correlate with pelvic pain scores in women with suspected endometriosis

The abundance of either CD14^high^ or CD14^low^ PMφ was compared to pain scores in women with suspected endometriosis and Pearson correlation calculated. No correlation was observed in women without endometriosis but this subgroup had the fewest patients (n=8; supplementary figure 2). When only women with diagnosed endometriosis (AFSI-IV) were assessed, neither CD14^high^ PMφ (Fig. A, p=0.058) nor CD14^low^ PMφ significantly correlated with pain scores (Fig. 4B, p=0.067), although a trend for negative and positive correlation, respectively, was evident. In women with suspected endometriosis (no endo and AFSI-IV combined), who were not receiving hormone treatment, CD14^high^ PMφ negatively correlated with pain scores (Fig. 4C, p=0.031) and CD14^low^ PMφ positively correlated with pain scores (Fig. 4D, p=0.032).

**Figure 4.**
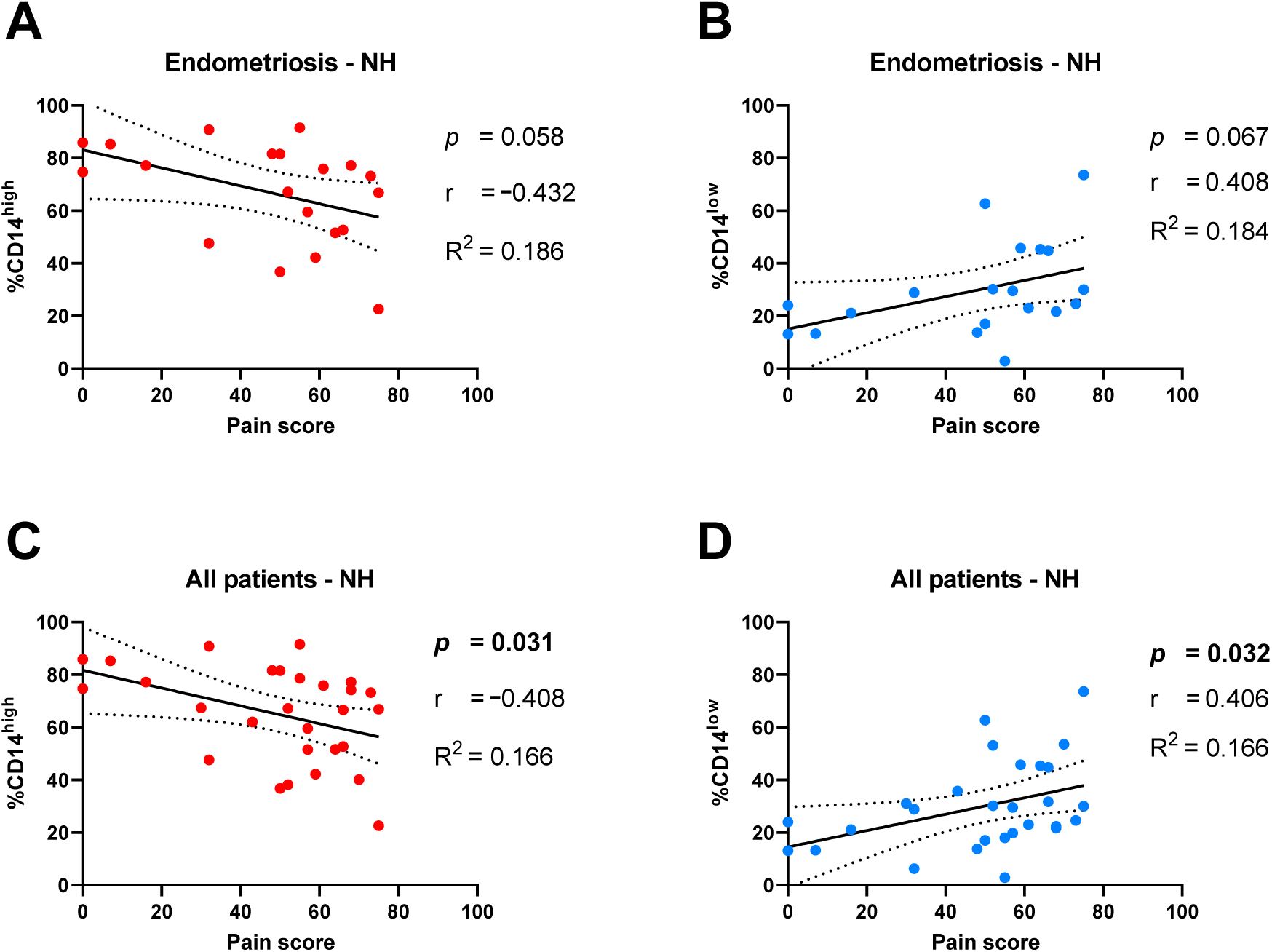
Peritoneal macrophages correlate with severity of pelvic pain in women with suspected endometriosis. **A** Frequency of CD14^high^ PMφ did not significantly correlate with pain scores in endometriosis patients not on hormones (n=21). **B** Frequency of CD14^low^ PMφ did not significantly correlate with pain scores in endometriosis patients not on hormones (n=21). **C** Frequency of CD14^high^ PMφ negatively correlated with pain scores in all patients with suspected endometriosis and not on hormones (p<0.05; n=28). **D** Frequency of CD14^low^ PMφ positively correlated with pain scores in all patients with suspected endometriosis and not on hormones (p<0.05; n=28). P value, correlation coefficient (r) and goodness of fit (R^2^) calculated by Pearson correlation. Line of best fit (black line) with 95% confidence intervals (dashed line). NH – not on hormones.

## Discussion

Peritoneal macrophages are believed to play a fundamental role in the pathophysiology of endometriosis (1, 2, 4, 24) but to date, information regarding different phenotypes of macrophage and how they relate to pain symptoms has been limited. In this study, we performed multi-parameter flow cytometry analysis of peritoneal fluid immune cells form women with suspected endometriosis and identified two populations of PMφ, CD14^high^ and CD14^low^. Most notably, we believe our data are the first to demonstrate that CD14^high^ PMφ are increased in women with laparoscopically-diagnosed endometriosis and that both CD14^high^ and CD14^low^ PMφ populations correlate with pain symptoms independent of endometriosis diagnosis.

We assessed pain scores in 54 women undergoing diagnostic laparoscopy for suspected endometriosis. Pelvic pain symptoms were recorded using the pain dimension of the EHP-30 questionnaire, an extensively validated endometriosis-specific health-related quality of life measurement (25, 26). We found that pain scores in women with endometriosis (mean and SEM, 45.5±5.54) were comparable to previous reports using the EHP-30 questionnaire in large cohorts of women from the UK (mean pain score 43.3, n=594 women), USA (mean pain score 49.3, n=225 women) and Australia (mean pain score 40.7, n=189 women) (27-29). These studies only assessed pain scores in women with surgically diagnosed endometriosis but in the current study we also assessed pain scores in women who did not receive a diagnosis of endometriosis at laparoscopy. Interestingly, mean pain score in the ‘no endo’ group was actually higher (mean and SEM, 57.3±4.78) than women with endometriosis but we found no significant difference between pain scores in women with or without endometriosis. Pain scores were not significantly different in patients graded as AFS stage I (47.3), II (38.3) or III/IV (47.8). Thus, in the current patient cohort, neither diagnosis of endometriosis nor AFS stage affected pain scores.

We profiled PMφ subpopulations in women with suspected endometriosis to determine if specific populations are altered by endometriosis stage, symptoms, or treatment. We identified two populations of peritoneal macrophage, CD14^high^ and CD14^low^, which, based on relative abundance and comparative transcriptional profiling in other studies (17), we believe to be equivalent to large (LpM) or small (SpM) peritoneal macrophage populations described in mice (30). Injection of endometrial tissue to induce lesion formation is associated with an increase in total peritoneal macrophages (4) and is associated with decreased LpM and increase SpM in mouse models of endometriosis (24). In our study, we detected an increase in CD14^high^ PMφ in women with endometriosis, consistent with previous reports assessing PMφ as single population in mouse and human (4, 8). However, the increase we detected was specific to the CD14^high^ PMφ subpopulation and is contrary to the decrease in the equivalent LpM population reported in the mouse model of endometriosis described by Yuan et al. (24). In response to peritoneal cavity inflammation in mice, LpM decrease and this is associated with infiltration of monocytes that replenish the LpM pool. Notably, it is not possible to distinguish between ‘resident’ PMφ and infiltrating monocytes in women as they both express CD14. Thus, the increase in CD14^high^ cells we detected in women with endometriosis may in part be accounted for by monocyte infiltration. This may also be reflected by the increase in the proportion of CD14^high^ relative to CD14^low^ we detected with AFS stages I, II and III/IV which may be associated with increased peritoneal inflammation in response to disease burden. Future studies will need to profile paired monocyte and PMφ samples from women with endometriosis in order to distinguish their respective contribution to the peritoneal environment in endometriosis.

Having identified two populations of peritoneal macrophage that were altered by presence of disease we next assessed whether these populations could offer insight into pain symptoms in endometriosis. A positive correlation was found between CD14^low^ PMφ and pelvic pain scores and a negative correlation between CD14^high^ PMφ and pelvic pain scores. These correlations were significant when all patients were assessed and were independent of endometriosis diagnosis. This may indicate that pelvic pain is more strongly associated with peritoneal macrophage dysfunction than the presence of active endometriosis lesions *per se*.

Peritoneal macrophages change throughout the inflammatory response, resulting in a state of adaptive homeostasis. Thus, although some patients may have no ‘active’ or visible disease at time of laparoscopy they may exhibit detectable changes in macrophages which correlate to pelvic pain. A previous study reported a correlation between concentrations of IL6 in peritoneal fluid and pelvic pain symptoms in women with endometriosis (31). Notably, PMφ are reported to be the principle source of IL-6 and this cytokine is increased in peritoneal fluid in women with endometriosis (32). Pain symptoms may therefore be affected by secretion of pro-inflammatory cytokines from PMφ in response to ‘active’ disease or as a legacy of previous inflammation from ‘historic’ disease. Thus, further profiling of peritoneal macrophages to account for both their phenotype and function could provide new insights into the relationship between endometriosis and pelvic pain symptoms.

Current treatment approaches for endometriosis focus on either surgical removal of disease or hormonal suppression of lesion growth but this does not improve pain symptoms in all women (22, 33, 34). Our new data suggest that divergence in pain responses may be influenced by the direct response of peritoneal macrophages to a given treatment. For example, GnRH agonists are reported to reduce pelvic pain in women with endometriosis (35) and, notably, this treatment has been shown to increase PMφ cytotoxicity (36) and reduce peritoneal fluid concentrations of the pain-correlated cytokine IL6 in women with endometriosis (37). Large-scale clinical trials have shown that the non-peptide GnRH antagonist Elagolix has a clinically meaningful reduction in the pain domain of EHP30 but only in around half of the women tested (22, 34). To the best of our knowledge, the impact of Elagolix on peritoneal macrophage function has not been investigated but baseline differences in peritoneal macrophage phenotype could account for some of the reported differences in pain response. We therefore propose that PMφ may have future utility for predicting response to treatment.

## Conclusions

The novel findings in this study provide new evidence for a link between abundance of peritoneal macrophage subpopulations and the experience of pelvic pain symptoms in women with suspected endometriosis. Collectively, our data suggest that the inflammatory profile of the peritoneal environment has a greater impact on pain symptoms than the presence/volume of visible disease. Reframing this perspective opens a path to future targeted therapies that aim to alter peritoneal macrophage function and reduce pelvic pain in women with endometriosis.

## Capsule

Peritoneal macrophage subpopulations are correlated with pelvic pain symptoms in women with suspected endometriosis and altered by presence of disease.

## Acknowledgements

We thank Dr Alistair Williams for histological evaluations of endometrial tissue and research team Helen Dewart, Jennifer Rowan and Ashleigh Symington for patient recruitment and sample collection. We thank members of PTKS laboratory for technical support, particularly Arantza Esnal-Zufiaurre for histology preparations and Olympia Kelepouri for sample processing. We thank Dr Luca Cassetta and Dr Samanta Mariani for advice setting up antibody panels. Flow cytometry data were generated with support from the QMRI Flow Cytometry and Cell Sorting Facility, University of Edinburgh. Studies were supported by MRC Programme Grants G1100356/1 to PTKS and MR/N024524/1 to PTKS, DAG and AWH. Studentship for BDL was funded by a grant to the MRC Centre for Reproductive Health (G1002033). AWH has received honoraria for consultancy for Ferring, Roche, Nordic Pharma, and Abbvie.

## Author contributions

Experimental design; DAG, AWH, PTKS, experimental procedures; DAG, BD, FC, manuscript preparation; DAG, FC, AWH, PTKS.

## Supplementary data

**Supplementary Table 1.** Age, BMI and EHP30 pain score of patients not on hormones did not significantly differ by presence/absence of disease or according to AFS stage – Variance between Endo and No Endo – Mann Whitney U test. Variance between AFS - Kruskal-Wallis test.

|  |  | No Endo | Endo | Endo by AFS stage |  |  |
| --- | --- | --- | --- | --- | --- | --- |
| AFS stage |  | N/A | I-IV | I | II | III/IV |
| N |  | 10 | 19 | 10 | 4 | 5 |
| Age | Mean | 29.1 | 31.83 | 29.67 | 33.5 | 34.4 |
|  | Std. Deviation | 6.28 | 7.09 | 7.583 | 5.447 | 7.403 |
|  | Std. Error of Mean | 1.986 | 1.671 | 2.528 | 2.723 | 3.311 |
|  | P value | 0.5308 |  | 0.4782 |  |  |
| BMI | Mean | 25.1 | 24.26 | 25.1 | 22 | 24.4 |
|  | Std. Deviation | 4.771 | 3.016 | 2.998 | 2.309 | 3.13 |
|  | Std. Error of Mean | 1.509 | 0.6918 | 0.9481 | 1.155 | 1.4 |
|  | P value | 0.6914 |  | 0.1876 |  |  |
| EHP30 pain | Mean | 57.33 | 45.53 | 47.3 | 38.25 | 47.8 |
|  | Std. Deviation | 14.35 | 24.14 | 24 | 30.58 | 23.59 |
|  | Std. Error of Mean | 4.784 | 5.537 | 7.591 | 15.29 | 10.55 |
|  | P value | 0.2576 |  | 0.741 |  |  |
| Infertility | (% True) | 20.0 | 42.1 | 30.0 | 75.0 | 40.0 |
| HMB | (% True) | 60.0 | 42.1 | 50.0 | 25.0 | 40.0 |

**Supplementary Table 2.** Age, BMI and EHP30 pain score of patients on hormones did not significantly differ according to AFS stage. When compared by presence/absence of disease ‘No Endo’ patients were younger. Variance between Endo and No Endo or AFS I vs AFS III/IV by Mann Whitney test.

|  |  | No Endo | Endo | Endo by AFS stage |  |  |
| --- | --- | --- | --- | --- | --- | --- |
| AFS stage |  | N/A | I-IV | I | II | III/IV |
| N |  | 10 | 10 | 8 | 0 | 2 |
| Age | Mean | 25.7 | 32.2 | 31.75 |  | 34 |
|  | Std. Deviation | 3.268 | 7.54 | 7.324 |  | 11.31 |
|  | Std. Error of Mean | 1.033 | 2.384 | 2.589 |  | 8 |
|  | P value | 0.049 |  | 0.6667 |  |  |
| BMI | Mean | 24 | 23.75 | 24.33 |  | 22 |
|  | Std. Deviation | 3.78 | 3.919 | 4.274 |  | 2.828 |
|  | Std. Error of Mean | 1.336 | 1.386 | 1.745 |  | 2 |
|  | P value | 0.8218 |  | 0.4286 |  |  |
| EHP30 pain | Mean | 57.13 | 53.89 | 50.14 |  | 67 |
|  | Std. Deviation | 13.94 | 15.85 | 16.07 |  | 4.243 |
|  | Std. Error of Mean | 4.93 | 5.282 | 6.073 |  | 3 |
|  | P value | 0.9074 |  | 0.1389 |  |  |
| Infertility | (% True) | 28.6 | 0.0 | 0.0 |  | 0.0 |
| HMB | (% True) | 33.3 | 55.6 | 42.9 |  | 100.0 |

**Supplementary Table 3.** Antibodies used in Flow cytometry analysis

| Antigen | Fluorophore | Cross- reactivity | Clone | Cat number | Supplier |
| --- | --- | --- | --- | --- | --- |
| CD3 | BV 711 | Human | OKT3 | 317327 | Biolegend |
| CD56 | BV 711 | Human | HCD56 | 318335 | Biolegend |
| CD19 | BV 711 | Human | HIB19 | 302245 | Biolegend |
| CD45 | PE-Texas red | Human | HI30 | MHCD4517 | Invitrogen |
| CD11b | BV 605 | Human | ICRF44 | 301331 | Biolegend |
| CD14 | BV 510 | Human | M5E2 | 301841 | Biolegend |
| HLA-DR | BV650 | Human | L243 | 307649 | Biolegend |

**Supplementary Figure 1.**
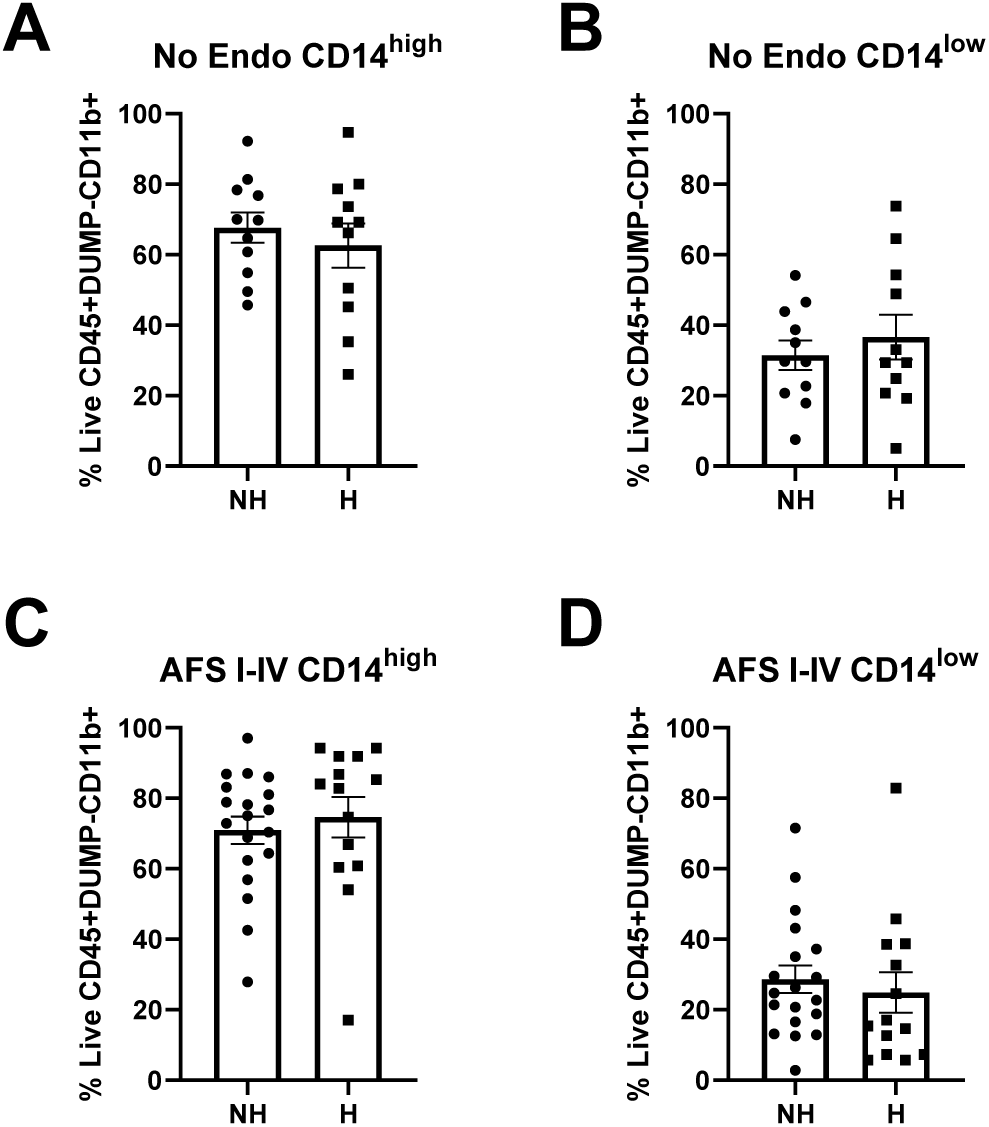
Peritoneal macrophages are not altered by hormone treatment. **A** Abundance of CD14 ^high^ peritoneal macrophages (PMφ) or **B** CD14 ^low^ PMφ in women without endometriosis was not altered by hormone treatment. **C** CD14 ^high^ and **D** CD14 ^low^ PMφ in women with endometriosis were not altered by hormone treatment. NH-No hormones, H - Hormones.

**Supplementary Figure 2.**
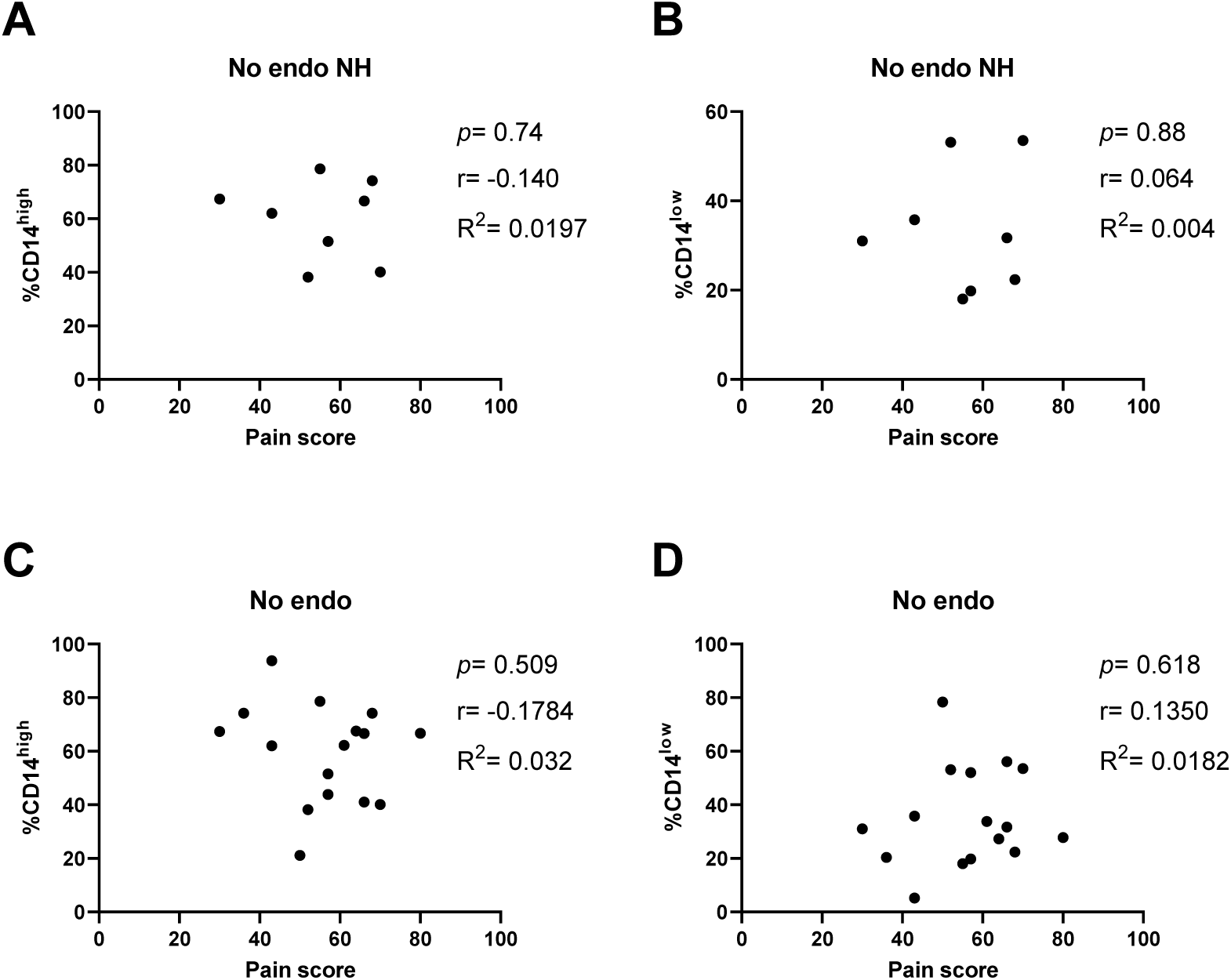
Peritoneal macrophages do not correlate with severity of pelvic pain in women without endometriosis. **A** Neither CD14^high^ nor **B** CD14^low^ PMφ correlated with pain scores in women without endometriosis and not on hormones (NH; n=8). **C** CD14^high^ or **D** CD14^low^ PMφ did not correlate with pain scores in women without endometriosis irrespective of hormone status (n=16).

